# Autoencoder-transformed transcriptome improves genotype-phenotype association studies

**DOI:** 10.1101/2023.07.23.550223

**Authors:** Qing Li, Jiayi Bian, Janith Weeraman, Albert Leung, Guotao Yang, Thierry Chekouo, Jun Yan, Jingjing Wu, Quan Long

**Author notes:** Corresponding authors; (**JW**) (**QL**). Equal contribution (Qing Li, Jiayi Bian).

## Abstract

Transcriptome-wide association study (TWAS) is an emerging model leveraging gene expressions to direct genotype-phenotype association mapping. A key component in TWAS is the prediction of gene expressions; and many statistical approaches have been developed along this line. However, a problem is that many genes have low expression heritability, limiting the performance of any predictive model. In this work, hypothesizing that appropriate denoising may improve the quality of expression data (including heritability), we propose AE-TWAS, which adds a transformation step before conducting standard TWAS. The transformation is composed of two steps by first splitting the whole transcriptome into co-expression networks (modules) and then using autoencoder (AE) to reconstruct the transcriptome data within each module. This transformation removes noise (including nonlinear ones) from the transcriptome data, paving the path for downstream TWAS. We showed two inspiring properties of AE-TWAS: (1) After transformation, the transcriptome data enjoy higher expression heritability at the low-heritability spectrum and possess higher connectivity within the modules. (2) The transferred transcriptome indeed enables better performance of TWAS; and moreover, the newly formed highly connected genes (i.e., hub genes) are more functionally relevant to diseases, evidenced by their functional annotations and overlap with TWAS hits.

## I. Introduction

The transcriptome-wide association study (TWAS) is a popular technique for integrating transcriptome data into genome-wide association studies (GWAS). Unlike GWAS, which associates phenotypic variance directly with genetic variants, TWAS quantitatively aggregates multiple genetic variants into a single test directed by transcriptome data. The key concept behind TWAS is genetically regulated expression (GReX), which is the component of gene expression attributed to genetic variants. The mainstream format of TWAS is a two-step protocol: First, using a reference dataset containing both genotype and expression data, a linear model is trained to predict GReX as a weighted linear combination of regulatory DNA elements. Techniques including ElasticNet [1], Bayesian sparse linear mixed models (BSLMM) [2] are used to train this genotype-based expression predictor. Then, in the second step, in the GWAS dataset which contains genotype and phenotype (but not expression), the above trained model is used to predict GReX which is subsequently associated with phenotypic traits. TWAS has since achieved popularity and success in identifying the genetic basis of complex traits [3]–[8], inspiring similar protocols for other endophenotypes such as IWAS [9], [10] for images and PWAS [11] for proteins.

Despite of its success in many projects, due to the low heritability of gene expressions, out of around 20,000 genes, usually only 7,000-10,000 genes may be analyzed in a TWAS project [1], [4], [12]. Although many efforts are devoted into the improvement of the predictions, current methods in TWAS [1], [8], [8], [13], [14] do not specifically handle the noise in the input gene expression profiles, including the influence of experimental artifacts, environmental factors, or, importantly, contributions from other interacting genes. Consequently, denoising data may improve expression heritability, which should enhance the genotype-expression model in the reference dataset. Although the current preprocessing methods can remove confounders such as PCs (principal components) in a linear model [15]–[17], the complex nonlinear confounders within the gene network are not addressed.

Towards this line, Autoencoder (AE) can denoise data and reduce complicated nonlinear confounders. AE is a specific type of neural network to encode input data into a compressed and informative representation (i.e., its latent variables), and then decode them back to reconstruct the input [18] (e.g., the configuration in **Fig. 1B**). By pursuing minimal discrepancies between the input and reconstructed, an AE effectively filters out biased noise from inputs and retaining essential patterns in the reconstructed, therefore lead to denoised outputs [19]. In cases where both the encoder and decoder employ linear operations, a linear AE is achieved, which provides latent variables equivalent to principal component analysis (PCA). Thus, AE can be considered as a nonlinear generalization of PCA, which has been used for processing expression data in a linear way [20].

**Fig. 1.**
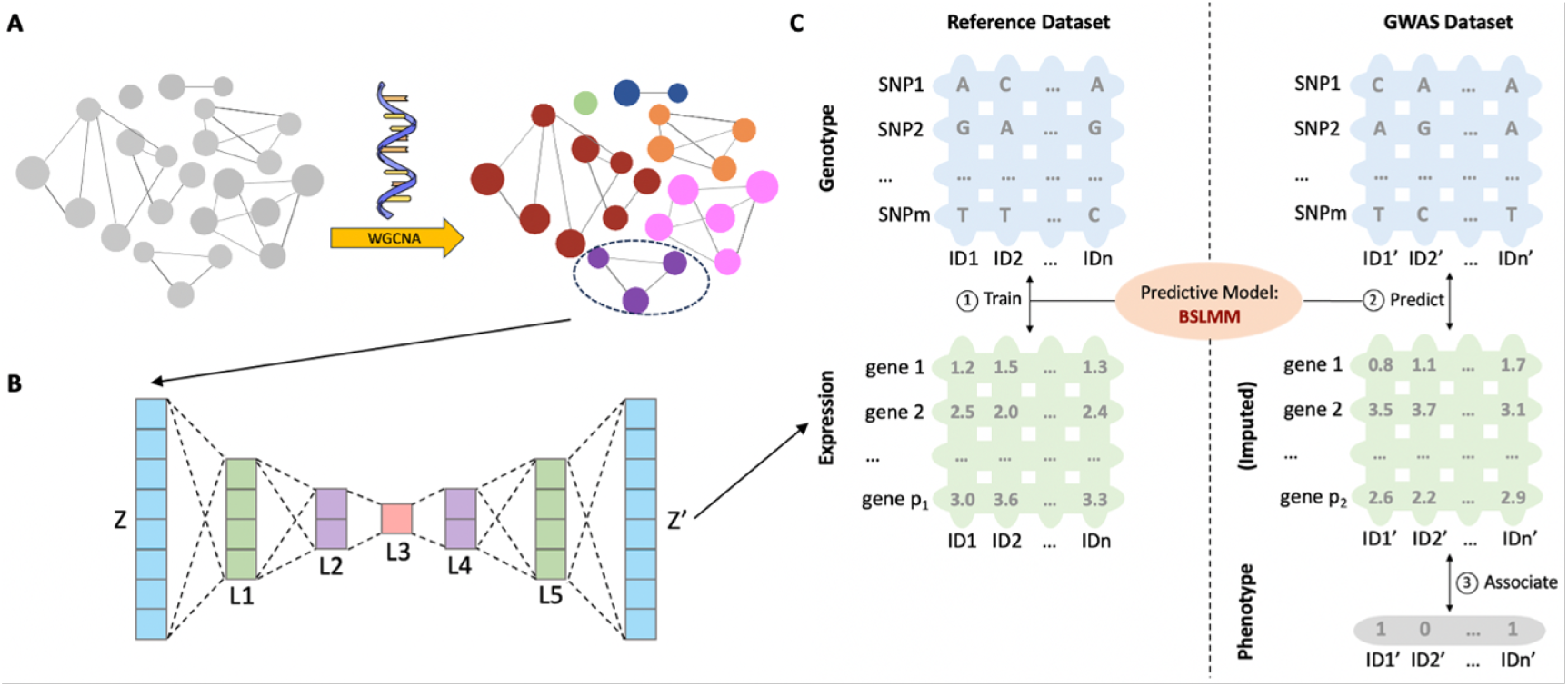
The AE-TWAS protocol. **A)** AE-TWAS first preprocesses transcriptome data by clustering genes into distinct modules using weighted gene co-expression network analysis (WGCNA). Each module contains a set of genes which are linked by substantial correlations (i.e., forming a co-expression network). **B)** AE-TWAS then applies a seven-layer conventional AE to the genes in each module. The AE outputs *Z*^*′*^ is referred to as AE-transformed transcriptome. **C)** AE-TWAS uses the transformed transcriptome *Z*^*′*^ to carry out TWAS. It employs the BSLMM to train the genotype-expression model in the reference dataset and then associates the predicted gene expressions to the traits in the target GWAS dataset.

To test the use of AE in data processing and its impact to the downstream expression-assisted association studies (i.e., TWAS), we implemented a new protocol, AutoEncoder-transformed TWAS (AE-TWAS), which utilizes AE-transformed transcriptome data into a TWAS protocol (**Fig. 1**). AE-TWAS first preprocesses transcriptome data by clustering genes into distinct modules using weighted gene co-expression network analysis (WGCNA) [21], [22]. Each module contains a set of genes which are linked by substantial correlations (i.e., forming a co-expression network) (**Fig. 1A**). Next, AE-TWAS applies a seven-layer conventional AE to the genes in each module (**Fig. 1B**). The AE outputs (*Z*^*′*^ in **Fig. 1B**) are referred to as AE-transformed transcriptome. Finally, AE-TWAS uses the transformed transcriptome to carry out TWAS (**Fig. 1C**). More specifically, AE-TWAS employs the widely used Bayesian sparse linear mixed model (BSLMM) [2] to train the genotype-expression model in the reference dataset and then associates the predicted gene expressions to the traits in the target GWAS dataset.

We applied AE-TWAS to whole-blood gene expression data on curated genes with 670 subjects in Genotype-Tissue Expression Project version 8 (GTEx v8) [16], [17]; and re-analyzed gene-trait associations on five diseases in three datasets, namely Wellcome Trust Case Control Consortium (WTCCC) [23], MSSNG [24] and Schizophrenia [25]. The applications yielded two unexpected and motivating discoveries. First, the heritability of most lowly heritable genes (*h*^2^ *<* 0.05) increased after AE transformation, although that for some high-heritability genes decreased (**Fig. 2**). Extending single genes to pairs of genes, the correlations between gene-pairs within co-expression networks increased significantly after the AE transformation (**Fig. 3**). Second, by comparing AE-TWAS identified genes to the ones identified by a standard TWAS protocol [1], it was observed that the AE-TWAS identified genes had higher proportions of genes that can be validated in DisGeNET, a comprehensive disease-gene database [26]– [29] in most diseases when compared to the standard TWAS protocol (**Fig. 4**). Surprisingly, such enhancement caused by AE transformation was also observed at the level of hub genes in the co-expression network (**Fig. 5**). Taking together, these results suggest that the AE-based denoising should be routinely adapted in expression data processing, just like the use of PCA currently, to facilitate many the downstream analyses including TWAS.

**Fig. 2.**
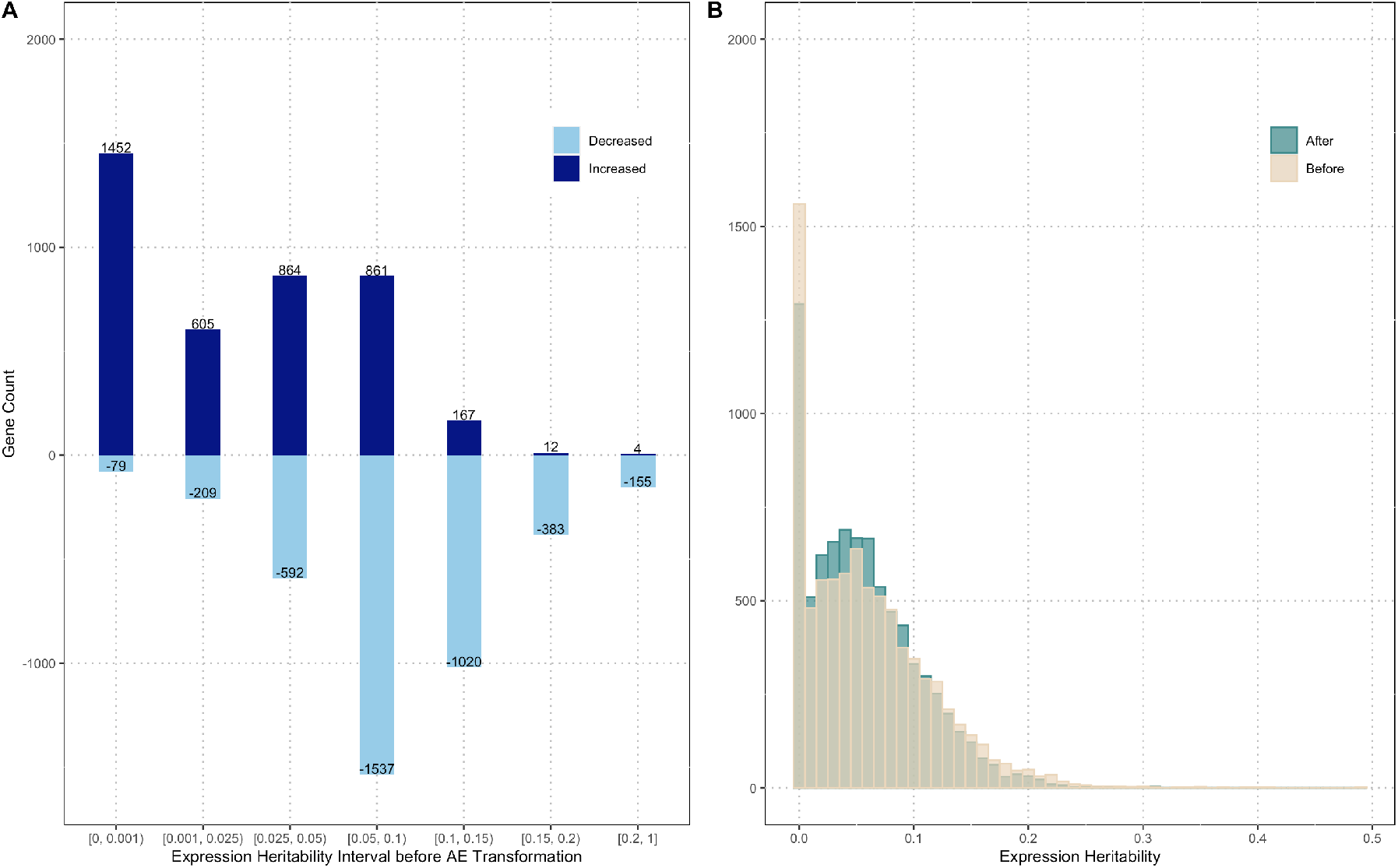
Expression heritability estimation of genes before and after AE transformation. **A)** Number of genes with increased and decreased expression heritability in each original expression heritability interval before AE transformation. **B)** Distribution of expression heritability for 8,191 genes before and after AE transformation.

**Fig. 3.**
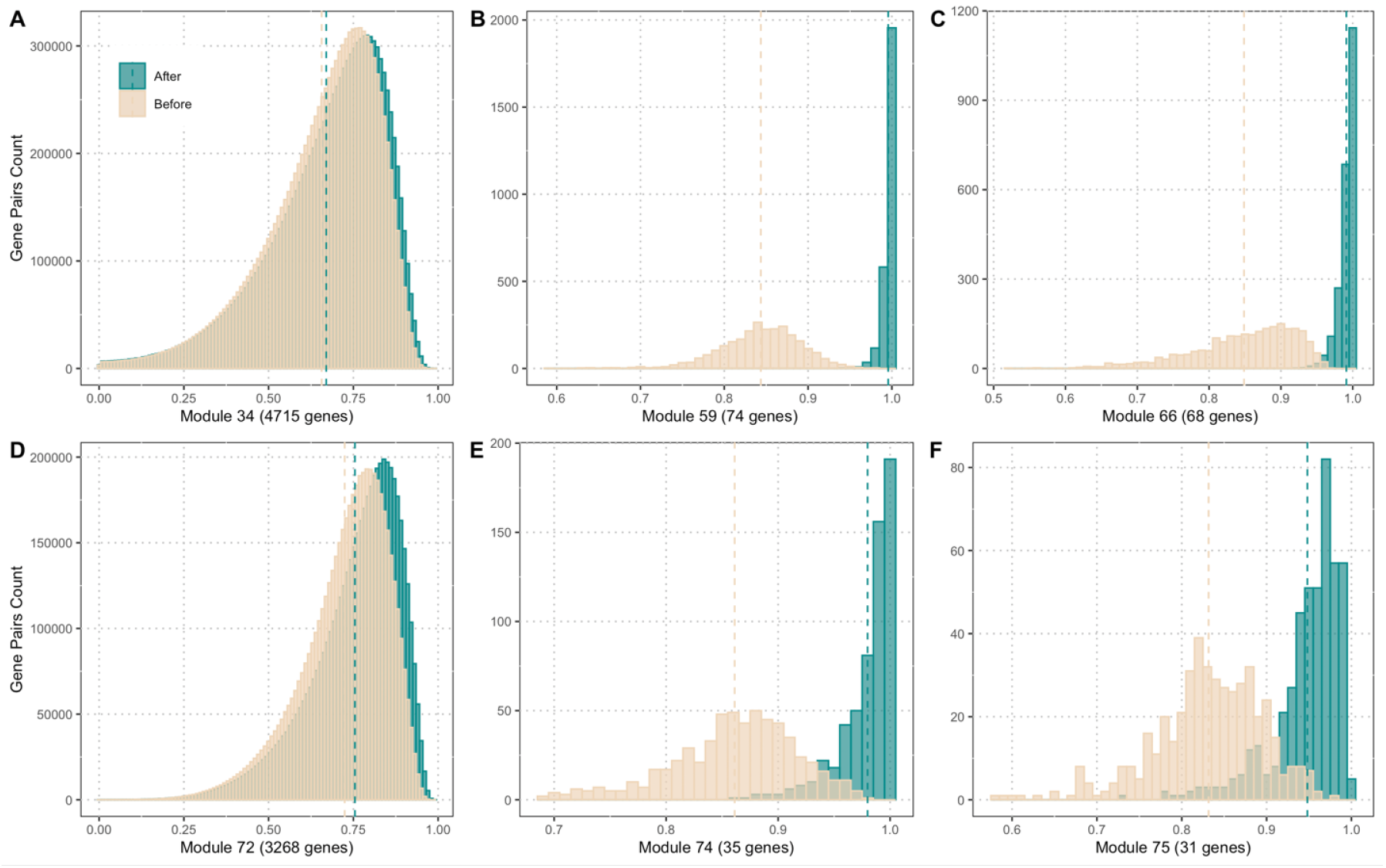
Histogram of absolute correlation among gene pairs within each module before and after AE transformation. Absolute correlation for each gene pair before AE transformation (yellow) and after AE transformation (green) are depicted. Each panel represents histogram for each module.

**Fig. 4.**
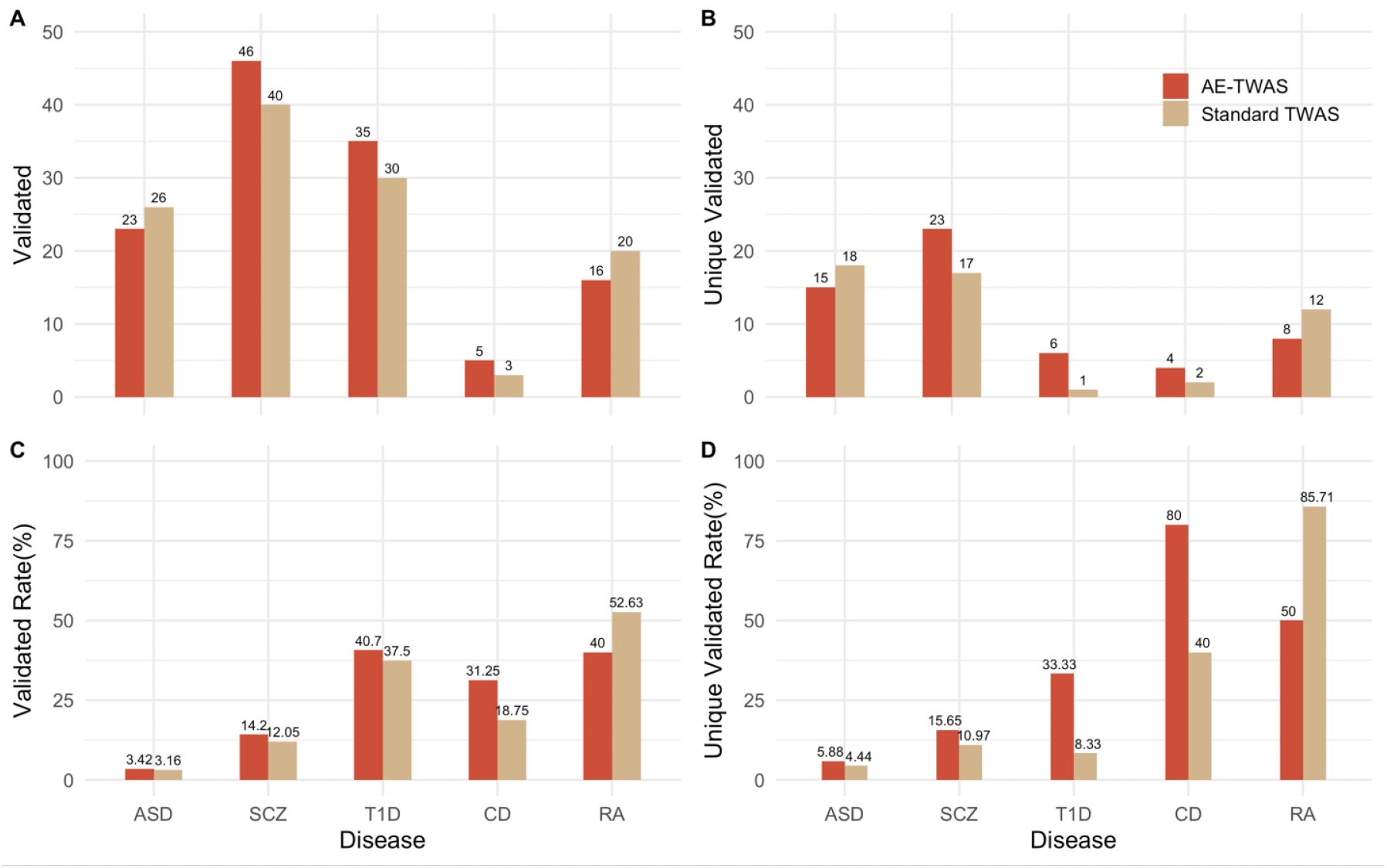
The DisGeNET functional annotations of genes identified by AE-TWAS and standard TWAS in five diseases.**A)** Number of validated genes (validated) reported in the DisGeNET database. **B)** Number of validated genes (unique validated) reported exclusively by one of the two protocols in the DisGeNET database. **C)** Proportion (validated rate) of all reported genes which are validated in the DisGeNET database. **D)** Proportion (unique validated rate) of genes reported exclusively by one of the two protocols in the DisGeNET database.

**Fig. 5.**
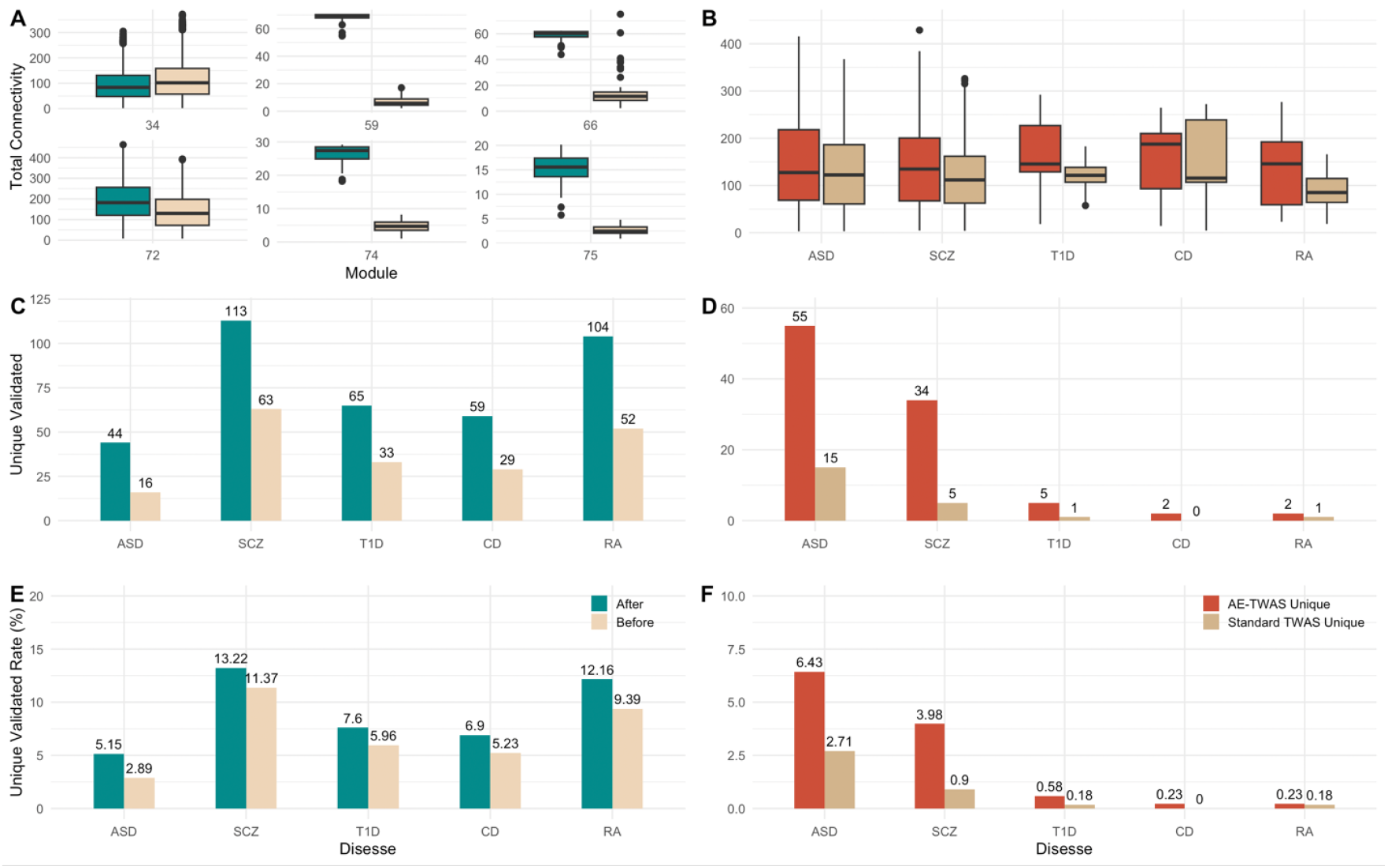
Network connectivity analysis of 8,191 genes clustered in 6 modules by WGCNA. **A)** Boxplot of total connectivity of 8191 genes within each module before and after AE transformation. **B)** Boxplot of total connectivity of uniquely discovered genes by AE-TWAS and standard TWAS in five diseases. **C)** Number of significant hub genes uniquely validated in DisGeNET for the five diseases before and after AE transformation. **D)** Number of significant hub genes uniquely validated in TWAS for the five diseases by the two protocols. **E)** Unique validation rate of significant hub genes validated in DisGeNET for the five diseases before and after AE transformation. **F)** Unique validation rate of significant hub genes validated in TWAS for the five diseases by the two protocols.

## II. Material and Methods

### A. Data sources and quality control

#### 1) Gene expressions

We obtained RNA-seq transcriptome data and genome-wide genotyping data from the GTEx v8 [16], [17]. We used whole-blood samples with 670 subjects, as this is the most representative general dataset when the disease-causing tissue is unknown. The target tissue initially contains 56,200 genes. After removing low-expression genes with average transcripts per million (TPM) below 1 and genes located outside of autosomes as we focused on human complex diseases unrelated to sex, 12,646 genes remained. Covariates, including genotyping principal components (PCs), were obtained from the GTEx portal. For each gene, we adjusted the gene expression for the top five genotyping PCs, age, sex and 60 confounding factors using a probabilistic estimation of expression residuals (PEER) analysis [15]–[17], following the GTEx best practices. In our analysis, we used log transformed gene expression data after covariate adjustment (*log*_2_(1+*gene expression*)) to maintain gene expression values in a reasonable scale.

#### 2) GWAS datasets

We used five diseases datasets (**Supplementary Table 1**). The WTCCC dataset includes approximately 2,000 individuals for each of seven diseases and a shared set of around 3,000 controls: type 2 diabetes (T2D), type 1 diabetes (T1D), hypertension (HT), rheumatoid arthritis (RA), coronary artery disease (CAD), Crohn’s disease (CD), and bipolar disorder (BD) [23]. We focused our analysis on T1D, CD, and RA, as few genes have been identified in association with the other diseases. The MSSNG database provides whole genome sequencing (WGS) and their phenotype data for over 11,500 individuals with autism spectrum disorder (ASD) [24]. The Schizophrenia dataset (dbGaP phs000473.v2.p2) provides whole exome sequences (WES) and their phenotype data for over 12,000 individuals from a population-based Swedish case-control cohort with schizophrenia (SCZ) [25]. For quality control, we first excluded genetic variants with a minor allele frequency (MAF) lower than 1%, as well as variants that deviated from Hardy-Weinberg equilibrium (HWE). We then mapped all genotype data in our GWAS datasets to the Genome Reference Consortium Human Build 38 (GRCh38) using the UCSC LiftOver tool [30] to ensure consistent genomic coordinates with genotypes in GTEx v8. This process resulted in a pruned set of 392,937 shared variants with GTEx v8 in the WTCCC dataset, 5,057,872 shared variants in the MSSNG dataset, and 401,515 shared variants in the Schizophrenia dataset carried forward in our analysis.

### B. The AE-TWAS protocol and its applications

#### 1) Weighted gene co-expression network analysis (WGCNA)

WGCNA is an R package consisting of a comprehensive collection of R functions for performing various aspects of weighted correlation network analysis [21], [22]. In our study, WGCNA is used to form co-expression networks (modules). It follows a step-by-step protocol for network construction and module detection. WGCNA first defines the weighted network by its adjacency matrix *A*, a symmetric matrix with entries ranging from 0 to 1, where the component *a*_*ij*_ indicates the connection strength between gene *i* and gene *j*. This adjacency matrix is calculated by raising the co-expression similarity component *s*_*ij*_ to a power: 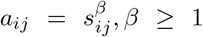, where *s*_*ij*_ is the absolute value of the correlation between gene *i* and gene *j*. The power *β* is selected using the function `pickSoftThreshold` in WGCNA based on the criterion of approximate scale-free topology.

Supported by WGCNA, AE-TWAS analyzed network topology on a set of candidate powers and chose the optimal power as the lowest at which the scale-free topology fit index reached 0.8. Using this criterion, the optimal power was set as 16. Afterward, AE-TWAS calculated the adjacency matrix A using the optimal power on gene expression data and transformed it into a topological overlap matrix (TOM) [31]. Modules were then detected through hierarchical clustering on the TOM using the dynamic tree cut method [32]. Finally, AE-TWAS merged modules with highly similar expression profiles based on their eigengenes, clustering them based on their correlation using a 0.75 cut-off. Genes that were not clustered into any module were removed for our subsequent analysis.

#### 2) Autoencoder transformation

AE-TWAS applied a seven-layer conventional AE to the genes in each module clustered by WGCNA in GTEx whole-blood samples. This conventional AE consists of one input layer, one output layer, and five hidden layers. Within each module, let *Z* represents the original input and *Z*^*′*^ is the reconstructed output. Let *q* be the number of nodes in the input and output layers, then the number of nodes in each of the five hidden layers are *q/*2, *q/*4, *q/*8, *q/*4, and *q/*2 respectively. Both the encoder and decoder in our seven-layer conventional AE are defined as fully connected multi-layer perceptron (MLP), and sigmoid functions were consistently used as activation functions. The optimal AE model is trained to minimize the reconstruction error with the least-square optimization: 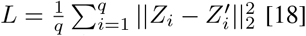.

To compare models’ performance, within each module, we divided the whole-blood samples randomly into training and testing sets using a 9:1 ratio, resulting in 603 subjects in the training dataset and 67 subjects in the testing dataset. We then normalized the data in both sets, scaling gene expression values for each gene to a range of (0,1) across all subjects. These normalized gene expressions values served as inputs of the AE models. We updated the weights using the adaptive moment estimation (ADAM) optimization algorithm [33]. The batch size was set equal to the size of the training set. We set the learning rate at 1*×*10^−3^ and the decay rate at 0.5 after 2,000 epochs, with a total of 20,000 epochs. The AE models were implemented using the PyTorch package.

#### 3) TWAS analysis based on Autoencoder-transformed transcriptomes

Assuming that the expression matrix *Z* is an *n×p*_1_ matrix where *n* and *p*_1_ represent the number of individuals and genes respectively. The transformed *Z*^*′*^ has the same dimensions of *Z*, and is utilized to train the predictive model in the reference dataset using BSLMM [2]:

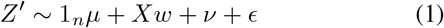

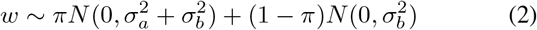

Here *X* is an *n×m* genotype matrix measured on the same *n* individuals at *m* cis-genetic variants (within 1 Mb of the gene), *w* is the corresponding effects of variants which come from a mixture of two normal distributions, ν is the random effect term, and *ϵ* is the error term.

AE-TWAS then predicted gene expressions as linear combinations of the effect estimates obtained from BSLMM and the genotypes derived from the GWAS dataset, which solely provides genotype and phenotype data. Finally, the imputed gene expressions, 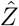, were associated with disease trait *Y* (an *n*^*′*^*×*1 vector where *n*^*′*^ represents the number of individuals in GWAS dataset) to identify significant gene-trait associations: 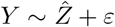.

We applied AE-TWAS on five diseases in three datasets (WTCCC, MSSNG and Schizophrenia datasets) [23]–[25] and compared their results in identifying significant gene-trait associations to a standard TWAS protocol [1]. Significant genes are defined as genes with p-values less or equal than 0.05 after false discovery rate (FDR) correction [34].

### C. Calculation of expression heritability

We estimated the gene expression heritability, which represents the proportion of expression variance explained by genotype, for genes in whole-blood samples using a variance-component model with a genetic relationship matrix (GRM) derived from genotype data. This estimation was conducted using the Genome-wide Complex Trait Analysis (GCTA) tool [35]. For each gene, variants located within 1 Mb of the gene’s start or end were used to construct the GRM. GCTA then calculates the proportion of the variance in gene expression attributed to these local variants using the following linear mixed model [36]:

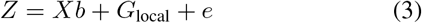

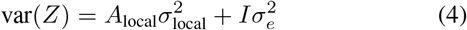

In this equation, *Z* represents gene expression and *b* is a vector of fixed effects. *A*_local_ refers to the GRM calculated from local variants, while the random effect *G*_local_ denotes the genetic effect attributable to the set of local variants, with var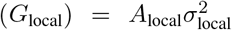. Variance components were calculated using restricted maximum likelihood (REML) [37].

### D. Calculation of gene pairs’ correlation and gene’s connectivity

We investigated the correlation among gene pairs within each module before and after AE transformation. The correlation between gene *i* and gene *j* is defined as cor(*x*_*i*_, *x*_*j*_) where *x*_*i*_ and *x*_*j*_ represent the expression profiles of the two genes. We considered the absolute value of the correlation. For 8,191 genes clustered in 6 modules, we calculated their total connectivity before and after AE transformation using `intramodularConnectivity` function in the WGCNA package [21], [22]. For each gene, the total connectivity is calculated as the sum of intramodular and extra-modular connectivity.

### E. Functional verification of discovered genes

To verify the functional relevance of gene-trait associations identified in the three GWAS datasets, we conducted a gene-disease search using the DisGeNET repository (v7.0), a platform containing 1,134,942 gene-disease associations (GDAs), between 21,671 genes and 30,170 traits [26]–[29]. For each disease, DisGeNET provides a specific summary of gene-disease associations, containing information on identified significant genes and their corresponding GDAs. These association summaries can be accessed using the Concept Unique Identifier (CUI) number assigned to each disease **Supplementary Table 1**).

## III. Results

### A. AE-transformation increased expression heritability and correlations between genes

#### 1) Seven-layer AE recovered transcriptome with high accuracy

The network construction and module detection protocol using WGCNA [21], [22](**Material and Methods**) on whole-blood gene expression data resulted in 6 modules encompassing 8,191 genes (**Supplementary Table 2**). The remaining 4,455 genes not clustered into any modules were excluded from subsequent analysis. We calculated the reconstruction accuracy of the optimal AE model within each module as the average *R*^2^ among all genes calculated from AE-transformed gene expressions and original normalized expressions, leading to high reconstruction accuracy ranging from 0.85 to 0.95 (**Supplementary Table 2**). These results indicate that the reconstruction accuracy is high, laying the ground for subsequent analyses.

#### 2) AE transformation improved expression heritability for lowly heritable genes

We evaluated the change in expression heritability for the 8,191 genes and visualized them based on their original expression heritability intervals before AE transformation. Among the seven expression heritability intervals, the low-spectrum intervals [0, 0.001), [0.001,0.025) and [0.025, 0.05) had a greater number of genes with increased expression heritability (**Fig. 2A**; **Supplementary Table 3**). This suggests that, after removing noises, the previously lowly heritable genes are now with higher expression heritability, allowing more genes to be TWAS-able. In contrast, the expression heritability of highly heritable genes reduced (**Fig. 2A**; **Supplementary Table 3**). We expected this as might be caused by the reduction of contribution from other genes. Such changes are also evident from the overall distributions of expression heritability before and after the transformation (**Fig. 2B**; **Supplementary Table 3**). Nevertheless, the current limitation of TWAS is largely due to that many genes have low expression heritability therefore cannot utilize expressions in the association mapping. The increase of expression heritability at this spectrum unlocks the potential of broader use of TWAS and other expression-directed association mappings.

#### 3) AE transformation increased correlation of gene pairs co-expression networks

Besides the expression heritability that is for single genes, we calculated the correlation (using absolute value) for all pairs of genes in each module before and after AE transformation (**Material and Methods**). Evidently, all modules experienced significant increase in absolute correlation among gene pairs following AE transformation (**Fig. 3**; **Supplementary Table 4**). It is noteworthy that all gene pairs (100%) within modules 59, 66, 74, and 75 showed a significant improvement in absolute correlation (*>* 0.1). Additionally, the majority of gene pairs in modules 34 (87.3%) and 72 (97.3%) exhibited an increase, although with a limited improvement in absolute correlation (0.01 for module 34 and 0.03 for module 72).

### B. AE-transformed expressions improved identification of gene-trait associations

To reveal the improvement of AE-transformed expressions, we carried out TWAS using expressions before and after transformation on five diseases including the MSSNG database for autism spectrum disorder (ASD) [24], a Schizophrenia dataset (SCZ) [25], and three WTCCC datasets [23], namely type 1 diabetes (T1D), Crohn’s disease (CD) and rheumatoid arthritis (RA) (**Material and Methods**). The full outcomes are elaborated in (**Supplementary Tables A**.**5-A**.**9**). We then examined the role of identified genes in each of the five diseases using DisGeNET [26]–[29], a well-established repository of validated gene-disease associations (**Material and Methods**).

### C. AE-transformed transcriptomes lead to TWAS outcomes with higher functional relevance

Using DisGeNET as the gold standard, we compared the number of validated associations and the validation rate (number of validated associations divided by the total number of discovered associations) (**Material and Methods**). After removing genes discovered by both methods, we also calculated the number of unique validated associations reported exclusively by one of the two methods and the unique validation rate. Looking at the numbers only, AE-TWAS model identified slightly more genes than the standard TWAS for T1D, CD, and RA in the WTCCC dataset, although did not outperform the standard TWAS on ASD (673 vs. 823) and SCZ (324 vs. 332) (**Fig. 4A**; **Supplementary Table 10**). The same trend was observed for unique validated associations (**Fig. 4B**; **Supplementary Table 10**). However, when considering the validation rate and unique validation rate, AE-TWAS outperformed the standard TWAS model in four of the five diseases with a large margin, with only RA being the exception. (**Fig. 4C and 4D**; **Supplementary Table 10**). Aggregately, the outcomes suggest that AE-TWAS outperformed the standard TWAS method in identifying gene-trait associations.

### D. AE-transformed genes enjoy higher co-expression network connectivity, which overlap with TWAS discoveries

We first checked the total connectivity of genes and learned that, except for module 34, the rest five modules enjoyed a significant increase of connectivity after AE transformation (**Fig. 5A**; **Supplementary Table 11**). The distributions of connectivity of uniquely discovered genes by AE-TWAS and standard TWAS protocols in five diseases also showed interesting properties: for all 5 diseases, the genes uniquely discovered by each protocol enjoyed increased connectivity after transformation (**Fig. 5B**; **Supplementary Table 12**).

### E. Hub genes in AE-transformed networks are more functionally relevant to diseases

To verify the functional relevance of the genes with high connectivity, we selected “hub genes” within each module using the cut-off of the 3^*rd*^ quantile of total connectivity within each module before AE transformation. This resulted in 2049 and 2350 hub genes, including 554 and 855 unique hub genes before and after AE transformation. For these hub genes, we then examined their functional roles in DisGeNET and in TWAS for the five diseases. Functional verification using DisGeNET showed that all the 5 diseases enjoyed increased number of validated and unique validated associations, as well as the validation rate and unique validation rate after AE transformation (**Fig. 5C and 5E**; **Supplementary Table 13**). We then further checked whether these hub genes are discovered by TWAS before transformation (or AE-TWAS after transformation, respectively) and verified by DisGeNET. Consistent to the above messages, all the 5 diseases enjoyed increased number of unique validated associations and unique validation rate after transformation (using AE-TWAS protocol) (**Fig. 5D and 5F**; **Supplementary Table 14**).

### F. Literature support of hub genes uniquely identified by AE-TWAS and validated in DisGeNET

We conducted a literature search on those genes that became hub genes after AE transformation and were both discovered uniquely by AE-TWAS and validated in DisGeNet. By conducting literature search of these genes, we were able to annotate the functional relevance between genes and related diseases. We annotated one gene for each of the five diseases that has the highest GDA score in DisGeNet [26]–[29] (**ASD: Supplementary Table 5**; **SCZ: Supplementary Table 6**; **T1D: Supplementary Table 7**; **CD: Supplementary Table 8**;**RA: Supplementary Table 9**). The gene names and associated p-values and scores are presented in **Table I**, where the genes for which we have conducted literature search below are colored in red.

**TABLE I.** Description of the selected hub genes in each of the five diseases in literature search.

| Gene Name | Disease | p-value from standard TWAS | p-value from AE-TWAS | DisGeNET score | PubMed IDs |
| --- | --- | --- | --- | --- | --- |
| *RNF135* | ASD | Not significant | 4.53E-02 | 0.10 | 30763456 |
| *FABP1* | ASD | Not significant | 3.55E-02 | 0.01 | 29754444 |
| *PSMB8-AS1* | T1D | Not significant | 3.07E-61 | 0.10 | 17632545 |
| *HEATR3* | CD | Not significant | 1.55E-06 | 0.01 | 23615072 |
| *SNX20* | CD | Not significant | 1.09E-04 | 0.11 | 27693347, 28008999, 17804789 |
| *PPP1R10* | RA | Not significant | 3.27E-02 | 0.10 | 19503088 |
| *IGHG1* | SCZ | Not significant | 3.96E-12 | 0.01 | 31837042 |
| *IGHG3* | SCZ | Not significant | 2.95E-12 | 0.01 | 30053866 |
| *SDCCAG8* | SCZ | Not significant | 6.16E-03 | 0.41 | 30285260, 31268507, 31374203 |
| *ASH2L* | SCZ | Not significant | 2.43E-02 | 0.20 | 23932495 |
| *NPTN* | SCZ | Not significant | 3.05E-02 | 0.32 | 27215477, 17123723 |
| *IFNGR2* | SCZ | Not significant | 2.31E-02 | 0.01 | 27563937 |
| *TWF2* | SCZ | Not significant | 3.69E-02 | 0.10 | 28540026 |

#### 1) Autism Spectrum Disorders (ASD)

[*RNF135*]: The ring finger protein 135 gene encodes an E3 ubiquitin ligase expressed in the cortex and cerebellum. *RNF135* was previously reported as one of the disease genes in individuals with ASD when studying the etiological heterogeneity of ASD [38]. Genetic analysis of the coding sequence of *RNF135* in a French cohort of patients with autism observed a significantly increased frequency of genotypes carrying the rare allele of the rs111902263 missense variant in patients, indicating the importance of the ubiquitin pathway in the aetiology of autism [39]. Moreover, McCallum et al. proposed new developments of empirical Bayes scan statistics and applied to a whole-exome sequencing study for ASD. They identified several promising candidate genes and *RNF135* was detected by all three scan statistic methods [40].

#### 2) Schizophrenia (SCZ)

[*SDCCAG8*]: The ciliopathy gene *SDCCAG8* encodes a centrosome associated protein. Genome editing of *SDCCAG8* caused defects in primary ciliogenesis and cilium-dependent cell signalling. Transcriptomic and phenotypic analysis of *SDCCAG8*-deficient cells showed that *SDCCAG8* influenced neurodevelopmental processes and is proved to be associated with SCZ [41]. Besides, *SDCCAG8* was identified as one of 7 SCZ risk genes that have centrosomal functions. Flynn et al. suggested that loss of *SDCCAG8* impairs cells’ ability to make primary cilia and that their capacity to repair genome damage may be reduced [42]. Moreover, gene enrichment analysis showing the enrichment of *SDCCAG8* as one of the cadherin genes in five major psychiatric disorders including SCZ [43].

#### 3) Type 1 Diabetes (T1D)

[*PSMB8-AS1*]: This gene is *PSMB8 antisense RNA 1* and we were able to annotate the functional relevance of *PSMB8* in T1D. *PSMB8* encodes a member of the proteasome B-type family, also known as the T1B family, that is a 20S core beta subunit. T1D is an autoimmune disease where local release of cytokines such as IL-1*β* and IFN-*γ* contributes to *β*-cell apoptosis. To identify relevant genes regulating this process, Lopes et al. performed a meta-analysis of 8 datasets of *β*-cell gene expression after exposure to IL-1*β* and IFN-*γ*. Genes were ranked according to their differential expression within and after 24 h from exposure and *PSMB8* was listed as one of the tops up-regulated genes [44]. Moreover, strongest known associations with T1D map to classical HLA class II genes. Sticht et al. performed a comprehensive HLA-wide genetic association analysis of T1D using whole-exome sequencing data in the large UK Biobank dataset. They applied gene-based association tests plus exon-based and single-variant association tests as complementary analysis. *PSMB8* was found to be significantly associated with T1D using the linear kernel SKAT test after Bonferroni correction for 1,209 exons [45].

#### 4) Crohn’s Disease (CD)

[*SNX20*]: *SNX20* interacts with the cytoplasmic domain of *PSGL1* and cycles *PSGL1* into endosomes. *SNX20* gene was added manually as one of the 126 significant genes associated with CD in a systematic review and functional analysis of a gene set potentially related to CD [46]. Besides, to increases the power of GWAS studies, Abegaz et al. studied the impact of confounding in gene–gene interaction (epistasis) detection. They introduced Model-Based Multifactor Dimensionality Reduction-Principal Component (MBMDR-PC) for Structured Populations and applied this method on International Inflammatory Bowel Disease Genetics Consortium Crohn’s disease (CD) data from 15 countries. *SNX20* was identified in 4 of the 8 pairs of potentially significant interacting genes which 36 of 109 interacting pairs of SNPs were mapped to [47].

#### 5) Rheumatoid Arthritis (RA)

[*PPP1R10*]: This gene encodes a protein phosphatase 1 binding protein. RA is an autoimmune disease results from a loss of tolerance to self-antigens in genetically susceptible individuals. Abegaz et al. reported a PhIP-Seq analysis of autoantibody repertoires from 64 RA patients, for comparison to a set of 73 healthy controls. A screen of synovial fluids and sera revealed novel disease-associated antibody specificities that were independent of seropositivity status. *PPP1R10* was detected as one of the significant disease-specific autoantibodies in Peptide/ORF enrichments associated with RA [48].

## IV. Conclusion

In summary, we presented the AE-TWAS framework, which integrates gene expressions after AE transformation into TWAS studies. We trained AE models on GTEx v8 using gene networks (modules) generated by WGCNA and conducted TWAS with BSLMM using gene expressions before and after transformation. We evaluated the change in expression heritability of genes and correlation between gene pairs within each module before and after AE transformation. Furthermore, we compared our framework to standard TWAS in identifying genes associated with five different diseases and characterized the functional relevance in relation to the transformation-altered connectivity and hub genes.

The impact of our study is beyond just to elucidate the genetic architecture of complex human diseases. It reveals that by harnessing modern machine learning techniques, one can denoise data to further genuine biological discoveries. Specifically, in this work, we employed autoencoders (AE) to investigate nonlinear interactions among gene expressions and eliminate noise. To our knowledge, our research provides the first evidence that AE-transformed gene expressions improve expression heritability estimation for low-heritability genes and enhance the correlation between genes in clustered co-expression networks (modules). These results suggest that AE models can learn patterns among genes and produce reconstructed outputs with essential information. Therefore, such AE-based data processing may be routinely used in daily expression analysis, similar to the current standard protocol of using PCA (the linear special case of an AE). In this work, the modules are generated purely by calculating correlations between genes using WGCNA. In the future, selected pathway information may be used to form modules of genes as the input to AE [49], [50].

We acknowledge that our study has certain limitations. First, our evaluation focused solely on the performance using the BSLMM method for GReX prediction. However, our AE-TWAS framework can also be adapted to other TWAS methods, such as ElasticNet implemented by PrediXcan and kernel methods articulated by ourselves [8], [10], [51], [52]. We chose BSLMM because our previous experiences show its supremacy to ElasticNet [8] and popularity in the field. Second, due to the complicated correlation structure within the transcriptome and the black-box and nonlinear nature of AE models, it is challenging to perform a simulation (in which the gold-standard is known) that can quantify the outperformance of AE-TWAS framework. We also haven’t provided closed-form mathematical derivations to quantify the improvement theoretically. To ease these limitations, future research could involve applying the AE-TWAS framework to various gene expression datasets and evaluating its performance across a range of TWAS methods. Third, in our real data analysis of five disease datasets, we demonstrated that our AE-TWAS framework outperformed the standard TWAS in terms of both the validation rate and the unique validation rate for four diseases: T1D, CD, ASD, and SCZ. However, this was not the case for RA, which may be attributable to potentially under-explored genetic basis. Finally, The AE-TWAS framework can be extended to other omics data and TWAS methods, making it a versatile tool in association studies for uncovering the genetic basis of complex human diseases. We again leave this to the future work.

## V. Data and Code availability

All the real datasets containing raw sequencing data used for this work are publicly available from their respective studies. GTEx gene expression data is available at https://gtexportal.org/home/datasets; GTEx whole genome sequencing data is available at https://www.ncbi.nlm.nih.gov/projects/gap/cgi-bin/study.cgi?studyid=phs000424.v9.p2; WTCCC dataset is available at https://www.wtccc.org.uk/; MSSNG database is available at https://research.mss.ng/ and Schizophrenia dataset is available at https://www.ncbi.nlm.nih.gov/projects/gap/cgi-bin/study.cgi?studyid=phs000473.v2.p2. The code of AE-TWAS is freely available on GitHub under the MIT license and can be found at https://github.com/theLongLab/AE-TWAS.

## VI. Acknowledgements

This work is supported by the New Frontiers in Research Fund [Exploration NFRFE-2018-00748 to Q.L.], the Alberta Innovates LevMax-Health Program Bridge Funds [222300769 to Q.L.], the Canada Foundation for Innovation [36605 to Q.L.], NSERC Discovery Grant [RGPIN-2018-04328 to J.W. and RGPIN-2017-04860 to Q.L.], and the Alberta Innovates Graduate Student Scholarships to J.B..

## VII. Author Contributions

(Q.L.= Quan Long; Q.L.2 = Qing Li; J.W.= Jingjing Wu; J.W.2=Janith Weeraman)

Conceptualization: Q.L.2, J.B., J.W., Q.L.

Methodology: Q.L.2, J.B., J.W., Q.L.;

Developed the tool: Q.L.2, J.B.

Funding, Resources & Supervision: T.C., J.Y., J.W., Q.L.

Formal analysis and investigation: Q.L.2, J.B., J.W.2, A.L., G.Y., Q.L.;

Manuscript writing: Q.L.2, J.B. Q.L., with contributions from all authors.

