## Supplementary Figures for "Autoencoder-transformed transcriptome improves genotype-phenotype association studies"

### S1 Text: Literature support of hub genes uniquely identified by AE-TWAS and validated in DisGeNET

**Overview**. We conducted a literature search on those genes that became hub genes after AE transformation and were both discovered uniquely by AE-TWAS and validated in DisGeNet. By conducting literature search of these genes, we were able to annotate the functional relevance between genes and related diseases. We annotated one gene for each of the five diseases that has the highest GDA score in DisGeNet (Piñero et al., 2015, 2017, 2020, 2021) (**ASD: S5 Table; SCZ: S6 Table; T1D: S7 Table; CD: S8 Table; RA: S9 Table**). The gene names and associated p-values and scores are presented in (**S15 Table**). The genes for which we have conducted literature search are colored in red.

**S15 Table: Description of the selected hub genes in each of the five diseases in literature search.**

| **Gene Name** | **Disease** | **p-value by**  **standard TWAS** | **p-value by**  **AE-TWAS** | **DisGeNET score** | **PubMed IDs** |
| --- | --- | --- | --- | --- | --- |
| *RNF135* | ASD | Not significant | 4.53E-02 | 0.1 | 30763456 |
| *FABP1* | ASD | Not significant | 3.55E-02 | 0.01 | 29754444 |
| *PSMB8-AS1* | T1D | Not significant | 3.07E-61 | 0.1 | 17632545 |
| *HEATR3* | CD | Not significant | 1.55E-06 | 0.01 | 23615072 |
| *SNX20* | CD | Not significant | 1.09E-04 | 0.11 | 27693347,28008999,17804789 |
| *PPP1R10* | RA | Not significant | 3.27E-02 | 0.1 | 19503088 |
| *IGHG1* | SCZ | Not significant | 3.96E-12 | 0.01 | 31837042 |
| *IGHG3* | SCZ | Not significant | 2.95E-12 | 0.01 | 30053866 |
| *SDCCAG8* | SCZ | Not significant | 6.16E-03 | 0.41 | 30285260,31268507,31374203 |
| *ASH2L* | SCZ | Not significant | 2.43E-02 | 0.2 | 23932495 |
| *NPTN* | SCZ | Not significant | 3.05E-02 | 0.32 | 27215477,17123723 |
| *IFNGR2* | SCZ | Not significant | 2.31E-02 | 0.01 | 27563937 |
| *TWF2* | SCZ | Not significant | 3.69E-02 | 0.1 | 28540026 |
